# 4-Phenylbutyrate Rescue in *GABRA1* Variants Associated with Developmental Epileptic Encephalopathies: from Cell and Mouse Models to Human

**DOI:** 10.64898/2026.05.20.724359

**Authors:** Ziang (Debbie) Song, Kirill Zavalin, Wangzhen Shen, Melissa B. DeLeeuw, Genevieve X. Hunn, Ria S. Eda, Li Ma, Robert Carson, Jing-Qiong Kang

**Author notes:** Academic Editor: Firstname Last-name. **Citation:** To be added by editorial staff during production.

## Abstract

Disease variants in *GABR* genes encoding γ-Aminobutyric acid type A receptor (GABA_A_R) subunits are major causes of developmental and epileptic encephalopathies (DEEs). There is no effective treatment for these DEEs although the GABA_A_R is a major target for antiseizure drugs. We previously identified the therapeutic effect of 4-phenyl-butyrate (PBA) in *Gabrg2*^+/Q390X^ knockin DEE mice and now test the effect of the drug in *GABRA1* variants that encode the α1 subunit. We used a multidisciplinary approach including *in silico* structural modeling, flow cytometry, patch clamp recordings and biochemistry in conjunction with differential tagging of the wild-type and the mutant alleles to evaluate the effect of PBA on rescue of GABA_A_R subunit expression, surface trafficking, and function *in vitro* in heterologous HEK293T cell model and *in vivo* in *Gabra1*^+/A322D^ mice. We found that both total and cell surface α1 expression was reduced when the variant α1 protein was present; suggesting reduced functional receptor on the cell membrane and synapse. Patch clamp recordings identified α1 variants reduced GABA-evoked current amplitude. *In silico* prediction indicated reduced protein stability for *GABRA1* variants indicated by negative ΔΔG values. PBA increased both total and surface expression of wildtype α1 and α1 variants; and improved expression of both wildtype and variant α1 alleles when these were co-expressed. Importantly, PBA also increased the GABA_A_R expression in the thalamus of the *Gabra1*^+/A322D^ mice. This study indicates that PBA is a promising treatment option for DEEs associated with *GABRA1* mutations. Our previous work has demonstrated that PBA improves proteostasis by enhancing expression of the wildtype allele, repairing the mutant allele, and reducing endoplasmic reticulum stress. Therefore, it can mitigate seizures and improve neurobehavioral phenotypes at behavioral levels. Based on this and our previous work on *GABRG2* and *SLC6A1* mutations, we propose that PBA holds promise as a common medicine for multiple genetic neurologic disorders that share the proteostasis pathology with a broad clinical application in DEEs.

## 1. Introduction

Pathogenic variants in genes involved in inhibitory neurotransmission comprise a substantial fraction of known monogenetic causes of neurologic disorders, including epilepsies and more severe developmental and epileptic encephalopathies[1,2]. Among these, variants in *GABR* genes that encode subunits of γ-aminobutyric acid (GABA) type A receptors (GABA_A_Rs) are particularly prominent. GABA_A_Rs mediate most synaptic inhibition and a significant portion of tonic inhibition in brain, and are essential for controlling neuronal excitability. GABA_A_Rs are ionotropic ligand-gated chloride channels comprised of five subunits with nineteen known isoforms. Most synaptic GABA_A_Rs are comprised of two α1-3, two β2/3, and one γ2 subunit. The α1β2/3γ2 combination is by far the most prevalent in brain, and *GABRA1, GABRB2, GABRB3*, and *GABRG2* genes encoding these subunits are the most frequently occurring *GABR* pathogenic variants[3]. Most known cases are dominant *de novo* mutations with a smaller proportion of familial cases[3].

In recent years, our lab has begun developing a promising new therapy for monogenetic epilepsies including *GABR* disorders using 4-phenylbutyrate (PBA), a small molecule with chemical chaperone and histone deacetylate inhibitor properties that is approved by FDA for treatment of urea cycle disorders[4,5]. In many cases, *GABR* mutations destabilize the subunit protein, impacting intracellular trafficking and/or interaction with partnering subunits, which affects GABA_A_R expression and functional properties[3]. PBA remedies this patho-mechanism two-fold by stabilizing mutant protein in mild cases where this is possible, or by assisting clearance of severely misfolded proteins and improving expression of functional wildtype allele and partnering subunits. Previously, we demonstrated this for *GABRG2*(Q390X) variant associated with Dravet Syndrome, which exhibits a severe trafficking pathology where mutant protein is retained in the endoplasmic reticulum and has a dominant negative effect on partnering subunits, significantly reducing GABA_A_R expression and synaptic inhibition, and causing seizures[6]. Treatment with PBA promoted clearance of the mutant protein and improved expression of the functional wildtype allele and partnering subunits in cell and mouse models, improving synaptic inhibition and significantly reducing the seizure burden[7]. Additionally, we have been successful in using PBA to treat *SLC6A1* disorders, which have a similar patho-mechanism. For these disorders, mutant variants in GABA transporter 1 are frequently mis-folded, resulting in reduced expression and transporter-mediated GABA uptake. Treatment with PBA improved intracellular trafficking and expression of the transporter, increased GABA uptake in cells and the *SLC6A1*(S295L) mouse model; and reduced the mouse seizure burden and neurobehavioral phenotypes[8-12]. These studies formed the basis for a clinical trial for PBA therapy, which reported seizure reduction or resolution in 70% of the *SLC6A1* patients tested (in submission)[13].

Clinical trials for *GABR* are also under way, and a significant need exists for translational studies evaluating efficacy of PBA for *GABR* variants, including subunits that have not yet been evaluated including *GABRA1*. In this study, we evaluate the effect *GABRA1* mutations have on GABA_A_R subunit stability and expression, and determine if PBA can restore these deficits. We focus on *GABRA1* variants I148F, R214C, R214H, M253T, T292S, F325L (Figure 1A; Table 1) obtained from the Cure GABAA variants patient organization. Additionally, we include the A322D variant associated with autosomal dominant juvenile myoclonic epilepsy. A322D is the first identified *GABRA1* variant and has been extensively characterized, displaying a misfolding/trafficking pathology with a modest dominant negative effect on partnering subunits[14-18]. Functional studies have previously also been done on R214C, R214H, T292S; R214C and R214H were identified in patients with Dravet syndrome and exhibit gating defects, showing diminished GABA_A_R currents and differing reports regarding expression[19-23]. The T292S variant is associated with severe developmental delay without epilepsy, and shows increased GABA sensitivity and normal expression, indicating a gain in GABA_A_R function[24]. No functional characterization has yet been done for I148F, F325 identified in a patient with epilepsy[25], and M253T identified in a West syndrome patient[26]. To predict the destabilizing effects these mutations have on the subunit protein, we use computational tools, measure expression of these variants along with the wildtype allele and partnering subunits in the HEK293 heterologous expression system and *Gabra1*^+/A322D^ mice. Then, we test if PBA treatment can improve expression deficits.

**Table 1.**
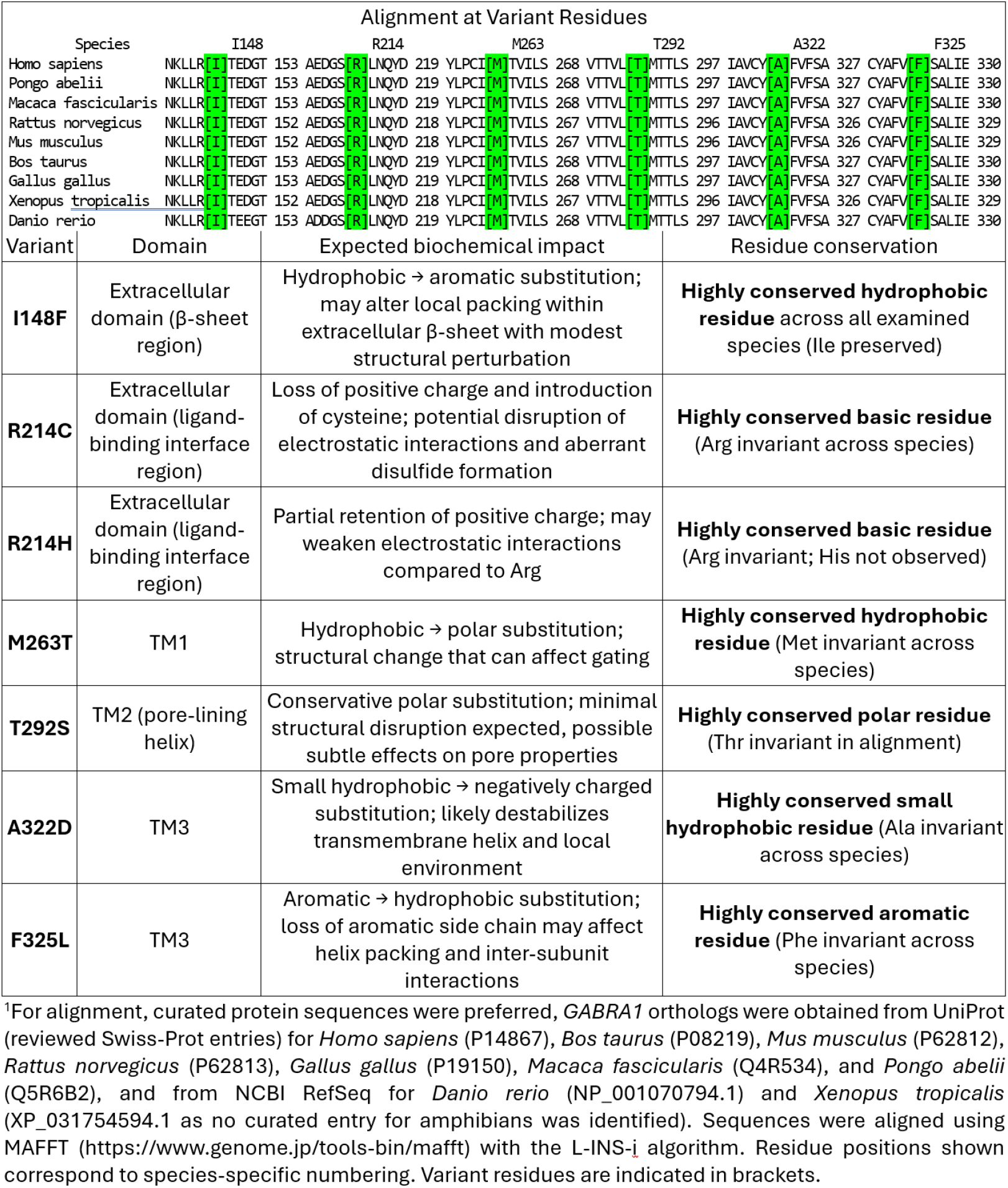
Alignment and putative mutation effects at *GABRA1* patient residues across vertebrates^1^.

**Figure 1.**
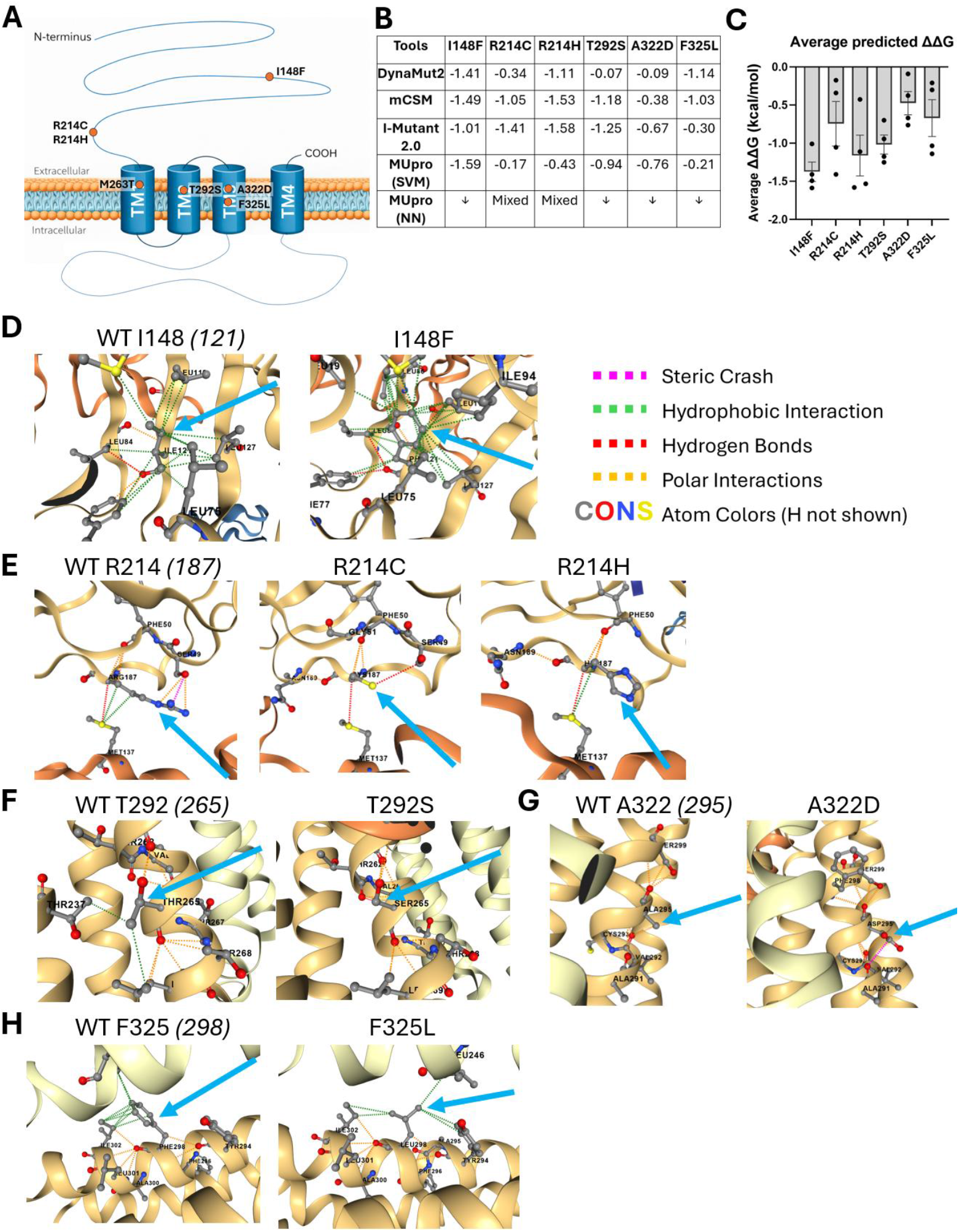
*GABRA1* variants disrupt local intermolecular interactions and destabilize the α1 subunit. (**A**) Location of variants included in this study within the α1 subunit. (**B**,**C**) Predicted changes in protein stability (ΔΔG) for selected *GABRA1* variants calculated using multiple *in silico* prediction tools. Table in (B) shows scores generated by each prediction tool, and bar graph in (C) shows the mean predicted ΔΔG for each variant. (**D**-**H**) DynaMut2-predicted local environments around the variant residues based on previously characterized GABA_A_R structure (PDB: 9CRS), chain D was used. For each variant, the WT structure is shown on the left and predicted structural change due to the variant is shown on the right. The peptide length used in the PDB: 9CRS structure report was slightly shorter than full length human α1. For each residue, the first peptide number corresponds to position within the full peptide, and the second number *(italicized in parentheses)* corresponds to the peptide number within the structure. The M263T variant was omitted due to inability to reliably predict its structure.

## 2. Materials and Methods

### Mice

The heterozygous *Gabra1*^*+/A322D*^ mouse line in C57BL/6J background was provided by Dr. Martin Gallagher at Vanderbilt University Medical Center. This line has previously been characterized, and genotyping was performed using Transnetyx using previously published probes[27]. Mice used in the study were males between 2 and 4 months old. Mice were housed in a temperature and humidity-controlled environment with a 12 hour light/dark schedule. Water and food was provided ad libitum. All animal procedures were performed in accordance with protocols approved by the Vanderbilt University Institutional Animal Care and Use Committee (IACUC). Animals were included based on confirmed genotype and normal health status. Animals with illness or unrelated injury were excluded. No collected animal data was excluded from analysis.

### GABA_A_R subunit cDNA and mutagenesis

Human α1, β2, and γ2S GABAA receptor subunit cDNAs were subcloned into the mammalian expression vector pcDNA3.1(+). An HA epitope tag was inserted into the extracellular N-terminus of the wild type α1 subunit of the mature peptide sequence. The HA tag was only added in the wild type α1 construct to allow selective detection of the wild type subunit protein in the heterozygous expression experiment. Site-directed mutations in the α1 subunit was generated using the Agilent QuikChange Site-Directed Mutagenesis Kit. All mutations were verified by Sanger DNA sequencing. Mutant plasmid DNA for transfection was prepared using the Qiagen MaxiPrep Kit.

### HEK293T cell transfection

Transient transfection was performed using polyethylenimine (PEI) in human embryonic kidney 293T (HEK293T) cells. HEK293T cells were seeded in 60mm dishes 24 hours prior to transfection. For each 60mm dish, 3 μg total plasmid DNA was transfected using PEI at a ratio of 2.5 μL PEI per 1 μg DNA. For homozygous receptor expression, α1, β2, and γ2S subunit cDNAs were transfected at a 1:1:1 ratio. For heterozygous α1 expression, equal amounts of wild-type and mutant α1 plasmids were co-transfected with β2 and γ2S cDNAs at a ratio of 0.5:0.5:1:1 for α1(mutant or control):α1^HA^(WT):β2:γ2S. After 48 hours of incubation, proteins are harvested or flow cytometry experiments were performed.

For electrophysiology experiments, a similar procedure was performed. HEK293T cells were transfected with 3 ug total plasmid DNA at a 1:1:0.5:0.3 ratio of α1:β2:γ2S:GFP using 5 μL Lipofectamine 2000 per 1 μg DNA; and cells were lifted using TrypLE to lift cells gently to preserve optimal cellular membrane conditions for patch clamp. Transfected cells were identified based on presence of GFP; typically, we observed 10-20% transfection rate.

### PBA administration *in vitro* and *in vivo*

The stock solution of PBA (Sigma-Aldrich, P21005) was prepared in dimethyl sulfoxide (DMSO). The optimal concentration (2 mmol·L^-1^) and treatment duration (24 h) for HEK293T cells were determined in a previous study and were used throughout this study. For *in vivo* experiments, mice received either vehicle or PBA (20 mg/mL in .9% saline, pH 7.4) at 100 mg/kg/day for 7 d through daily intraperitoneal injections; this dose was determined based on our prior experiments in *Slc6a1*^+/S295L^ mice[9] and is comparable with 12.4 g/m^2^/day dosing used in clinical trial patients[13]. Brain tissues were collected 24 hours after the final PBA administration (day 8) for subsequent Western blot analysis. To minimize potential confounders, treated and untreated animals were derived from the same litters and treated with vehicle or PBA under identical conditions based on random assignment. Tissues were collected at the same time, and all samples were processed and analyzed in parallel. No formal a priori sample size calculation was performed; sample size was determined by the number of available mice.

### Surface expression of α1 subunit by flow cytometry

Protocol was based on our earlier studies[28,29]. Cells were detached by trypsinization, washed, and resuspended in FACS buffer (phosphate-buffered saline supplemented with 2% fetal bovine serum and 0.05% sodium azide). For surface expression analysis, non-permeabilized cells were incubated with anti-GABA_A_R α1 antibody for 2 hours and then incubated with fluorophore Alexa-647 conjugated goat anti-mouse secondary antibody (1:2000) for 1 hour at 4 °C. Cells were subsequently washed and analyzed on a 3-laser LSR II flow cytometer (BD Biosciences) at the Vanderbilt Flow Cytometry Core. Cell populations were first gated based on forward and side scatter (FSC/SSC) to exclude debris. Approximately 10,000 events were collected per sample. Untransfected cells and isotype controls were included to define background fluorescence. Surface α1 expression was quantified as mean fluorescence intensity (MFI) using FlowJo v7.1 (Tree Star, OR).

### Western blot

Protocol was based on our earlier studies[30,31]. The protocol was similar for mouse samples and HEK293T cells. Live transfected HEK293T cells were washed three times with phosphate-buffered saline (PBS, 1×, pH 7.4); alternatively, mouse brain samples were dissected in PBS. Then, samples were lysed in RIPA buffer (20 mM Tris, 20 mM EGTA, 1 mM DTT, and 1 mM benzamidine) supplemented with protease inhibitors (0.01 mM PMSF, 0.005 μg/mL leupeptin, and 0.005 μg/mL pepstatin) for 30 min at 4 °C. Cell lysates were sonicated and clarified by centrifugation. Protein concentrations were determined using the Bradford assay. Equal amounts of protein were mixed with Laemmli sample buffer, and separated by SDS-PAGE. Proteins were transferred onto PVDF membranes. Membranes were incubated overnight at 4 °C with primary antibodies. Primary antibodies used were mouse anti-GABAAR α1 1:500 (Millipore Sigma MABN489), rabbit anti-GABAAR β2 1:500 (Millipore Sigma AB5561), rabbit GABAAR β3 1:500 (Novus NB300-199), rabbit anti-GABAAR γ2 1:500 (Synaptic Systems 224003), rabbit anti-HA 1:500 (Cell Signaling Technology 3724), and anti-ATPase 1:1000 (Developmental Studies Hybridoma Bank a6F). Secondary antibodies were LI-COR IRDye 680LT Goat anti-Mouse IgG Secondary Antibody (926-68020) and IRDye 800CW Goat anti-Rabbit IgG Secondary Antibody (926-32211), both 1:10000. Band intensities were quantified using LiCor Image Studio software.

### Electrophysiology

Neurons were patched with 3-5 MΩ borosilicate glass electrodes (World Precision Instruments, TW150-4, pulled with Narishige PP-830 or Sutter P-2000 pullers) filled with internal solution containing (mM): 153 KCl, 10 HEPES, 1 MgCl_2_, 5 EGTA, and 1 MgATP (300 mOsm, pH 7.4) and clamped at -70 mV; external solution contained (mM): 142 NaCl, 8 KCl, 10 HEPES, 10 Glucose, 6 MgCl_2_, 1 CaCl_2_ (325 mOsm, pH 7.4). Whole-cell recordings were acquired using an Axon MultiClamp 700B amplifier, filtered at 2 kHz, digitized at 10 kHz with Digidata 1440A, and recorded with Clampex 10.4 software (Molecular Devices). Currents were elicited with 1 mM GABA application through a microcapillary using a gravity system. Peak current analysis was performed using ClampFit software.

### Predictive structural modeling

We simulated the perturbations of six α1 mutations with multiple AI modeling tools. The crystal structure of native human GABA_A_R (PDB ID: 9CRS) was retrieved from the RCSB Protein Data Bank. This structure was selected for its high resolution (2.9 Å) and complete β2-α1-β2-α1-γ2 subunit assembly. All six point mutations were introduced *in silico*, and DynaMut2 was used to visualize tertiary structural changes and to predict ΔΔG and residue-level flexibility based on normal mode analysis.

Structural stability changes induced by variants were predicted using the α1 subunit from chain D of the receptor model. Because α1 subunits occupy distinct positions within the pentameric assembly, the local structural environment varies slightly between chains due to differences in inter-subunit interactions, which can lead to modest variation in predicted ΔΔG values. For consistency, all variants were modeled using chain D.

### Data analysis and statistics

Statistical data was stored using Microsoft Excel, analyzed and graphed using GraphPad Prism. Final figures were made in Microsoft Powerpoint. Statistical tests are specified in each figure legend. Data were assumed to be normally distributed. Values are displayed as mean ± SEM with one-way ANOVA with Sidak’s or Dunnett’s posthoc multiple comparisons tests as appropriate. For Western blot experiments, values per blot were typically normalized to loading control, then to untreated WT sample within that blot, as specified in figures.

## 3. Results

### 3.1. GABRA1 variants destabilize α1 subunit structure and disrupt local intermolecular interactions

We began by performing *in silico* structural analysis of the mutations. GABA_A_Rs belong to a “cys-loop” superfamily of pentameric ligand-gated ion channels, and each subunit contains an extracellular N-terminal domain, four transmembrane (TM) domains with TM2 lining the channel pore, and an intracellular cytoplasmic loop between TM3 and TM4[32]. The full length of human α1 peptide is 456 amino acids. I148, R214 residues are in the extracellular domain: I148 in the β-sheet region and R214 in the ligand-binding interface region. M263, T292, A322, and F325 residues are in TM domains: M263 in TM1, T292 in TM2, and A322 and F325 in TM3 (Figure 1A). We performed sequence alignment and evaluated sequence conservation for these residues, for which we included *GABRA1* sequences from fish, amphibians, birds, and mammals (Table 1). All six residues were highly conserved across the species as were the neighboring residues, indicating their importance to GABA_A_R function.

We predicted structural implications of the variants using DynaMut2 with the human α1β2γ2 GABA_A_R structure (PDB: 9CRS), finding that most variants are particularly disruptive to native intermolecular interactions, including hydrogen bonds, polar contacts, and hydrophobic interactions (Figure 1B,C). R214 substitutions (R214C and R214H) alter charge and side-chain interactions, leading to rearrangement of local environment and destabilized contacts. The A322D variant introduces a negatively charged residue into a hydrophobic region, resulting in unfavorable polarity interactions and steric strain. The T292S substitution alters hydrogen bonding and local packing interactions despite being chemically conservative. In contrast, the F325L substitution preserves hydrophobic character with only minor packing changes, consistent with its comparatively mild destabilizing effect. The M263T variant was omitted due to inability to reliably predict structure.

We then estimated the change in protein stability for each mutation using an array of computational platforms: DynaMut2, mCSM, I-Mutant 2.0, and MUpro. After independently analyzing each variant within each platform (Figure 1B), we averaged the predicted changes in Gibbs free energy (ΔΔG) across the platforms for each variant (Figure 1C). Averaged ΔΔG values were consistently negative across all variants analyzed, indicating a destabilizing effect of the mutations: I148F (-1.375 kcal/mol), R214C (-0.744 kcal/mol), R214H (-1.162 kcal/mol), T292S (-1.018 kcal/mol), F325L(-0.671kcal/mol), A322D(-0.475kcal/mol).

Across prediction tools, we broadly ranked variants by magnitude of destabilization. R214H and I148F exhibited the strongest predicted destabilizing effects, with ΔΔG values frequently exceeding −1.0 kcal/mol across multiple algorithms. T292S and R214C showed moderate destabilization, while A322D displayed variable but overall destabilizing effects depending on the method used. F325L consistently showed the mildest destabilization, with smaller magnitude ΔΔG values and greater variability across tools. Given the destabilizing effect of the mutations, we concluded that these variants are likely to exhibit a misfolding and trafficking pathology that affects GABA_A_R expression and could be targeted by PBA therapy.

### 3.2. Most GABRA1 variants show reduced α1 expression in α1β2γ2-expressing HEK293T cells

Next, we evaluated the impact of *GABRA1* variants on α1 subunit expression using homozygous expression in the heterologous HEK293T cell system. We chose the α1β2γ2 as it is one of the most common synaptic GABA_A_R combinations and was consistent with our structural modeling. We expressed either the wild type (WT) or variant α1 co-transfected with β2 and γ2 subunits at a 1:1:1 ratio and examined α1 expression by Western blot (Figure 2A). We observed that I148F, R214C, R214H, M263T, T292S, and A322D variants all displayed a severe reduction in expression compared to WT (Figure 2B): 75.8% decrease for I148F (####P < 0.0001 vs WT), 72.0% for R214C (####P < 0.0001), 73.0% for R214H (####P < 0.0001), 74.5% for T292S (####P < 0.0001); M263T was more mild (61.5% reduction; ####P < 0.0001) and A322D was most strongly affected (85.2% reduction, ####P < 0.0001). F325L was an exception and showed expression comparable to WT (9.1% decrease, P = 0.3181).

**Figure 2.**
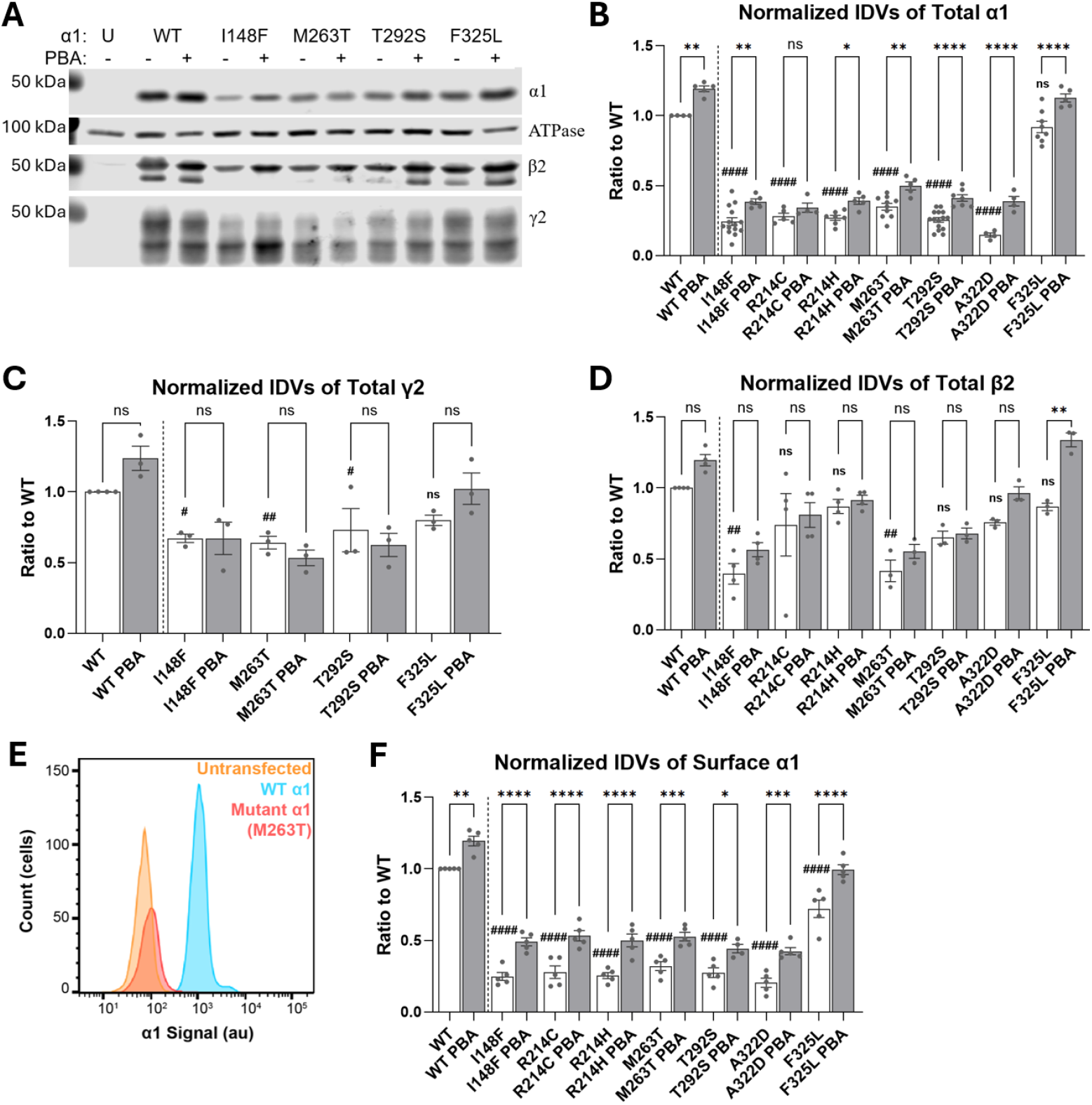
*GABRA1* variants show reduced α1 subunit expression that is partially rescued by PBA. (**A**) Western blot showing total expression of α1, β2, γ2 subunits in HEK293T cells cotransfected with α1, β2, and γ2S cDNAs (1:1:1 ratio). For each condition, we varied whether the WT or variant α1 was transfected; for PBA treatment, 2mM PBA was added 24 hours prior to harvesting. ATPase was used as a loading control. (**B**) Quantification of total α1 protein levels normalized to WT. (**C**-**D**) Quantification of total γ2 and β2 protein; not every α1 blot included a γ2 and β2 co-stain, which is reflected in the smaller sample number for these blots. (**E**-**F**) Surface expression of α1 quantified by flow cytometry using an anti-α1 antibody with Alexa-647 secondary. A representative example of flow cytometry results is shown in (E) with quantification normalized to untreated WT in (F). Data are presented as mean ± SEM with one-way ANOVA; ####, ##, #P < 0.0001, 0.01, 0.05 untreated variant vs untreated WT by Dunnett’s multiple comparisons test; ****, ***, **, *P < 0.0001, 0.001, 0.01, 0.05 PBA vs corresponding untreated by Sidak’s multiple comparisons test.

We also evaluated expression of the partnering β2 and γ2 subunits. For γ2, we were only able to evaluate expression for a subset of the variants I148F, M263T, T292S, and F325L. Compared to WT, γ2 expression was 32.9% lower for I148F (#P < 0.05), 36.0% for M263T (##P < 0.01), and 27.1% lower for T292S (#P < 0.05); and comparable to WT for F325L (Figure 2C). β2 expression was 40.5% lower that WT for I148F (##P < 0.01) and 58.5% lower for M263T (##P < 0.01; Figure 2D). We did not detect any statistically significant changes in β2 expression for R214C, R214H, T292S, A322D, and F325L.

### 3.3. PBA improves α1 expression for most GABRA1 variants

We next assessed whether α1 expression could be restored by treatment with PBA. We used a treatment protocol that we previously optimized to rescue expression of other *GABR* variants in HEK293T cells, which entails a 24 hour treatment with 2 mM PBA prior to cell harvest. PBA treatment significantly increased α1 expression for most variants (Figure 2B): 59.4% increase for I148F (**P < 0.01 vs untreated), 44.4% for R214H (*P < 0.05 vs untreated), 42.7% for M263T (**P < 0.01 vs untreated), 61.5% for T292S (****P < 0.0001 vs untreated). For R214C, treated expression values were 22.4% greater than untreated, but this increase was not statistically significant (P = 0.8350). A 19.2% increase was also observed for WT α1 (**P < 0.01 vs untreated); and 19.1% increase was observed for F325L (****P < 0.0001 vs untreated), which did not show a baseline reduction in expression. However, PBA treatment did not fully restore expression of α1 variants to wildtype levels; despite the improvements, all α1 variants that showed reduced expression at baseline remained at 34-50% of baseline WT levels following PBA treatment.

We detected no changes in γ2 for PBA vs untreated for WT, I148F, M263T, T292S, and F325L variants (Figure 2C); nor in β2 for WT, I148F, R214C, R214H, M263T, and T292S (Figure 2D). For F325L, β2 expression increased by 54.5% (**P < 0.01 vs untreated). Notably for WT and several variants, we noticed an upward trend in expression values, and it’s possible that a small percent increase could be resolved with higher statistical power.

### 3.4. PBA improves surface expression in α1β2γ2-expressing HEK293T cells

Next, we investigated α1 surface expression of the α1 variants using flow cytometry. Variants that showed reduced total α1 expression also showed a comparable reduction in α1 surface expression (Figure 2E-F): 75.2% reduction compared to WT for I148F (####P<0.0001), 72.2% for R214C (####P<0.0001), 74.6% for R214H (####P<0.0001), 68.0% for M263T (####P<0.0001), 72.6% for T292S (####P<0.0001), and 79.6% for A322D (####P<0.0001). F325L variant showed a moderate reduction in surface α1 expression (28.0% less than WT; ####P<0.0001) though total expression was not significantly reduced.

PBA treatment improved α1 surface expression for WT and all of the variants (Figure 2F): 19.4% increase vs untreated for WT (**P<0.01), 97.6% for I148F (****P<0.0001), 91.4% for R214C (****P<0.0001), 96.9% for R214H (****P<0.0001), 64.4% for M263T (***P<0.001), 61.5% for T292S (*P<0.05), 108% for A322D (***P<0.001), and 38.1% for F325L (****P<0.0001).

Additionally, we performed a limited electrophysiologic evaluation of whole-cell currents elicited by 1 mM GABA application. We evaluated WT and R214C, R214H, A322D, and F325L variants with and without PBA in HEK293T cells. The average amplitude of GABA-elicited currents was similar across the variants and appeared lower than WT (Figure S1A). However, we were unable to adequately appraise significance of individual variants due to low statistical power caused by high variance and low sample number. Therefore, we made a comparison across the similar values of pooled R214C, R214H, A322D, and F325L variants vs WT, which was significantly lower (10 vs 6.4 nA, *P < 0.05; Figure S1B).

### 3.5. Total and WT α1 expression with “heterozygous” α1α1^HA^β2γ2 expression

Next, we sought to recapitulate the heterozygous state of the patients by co-expressing α1 variants/WT α1 control together with an HA-tag labeled WT α1 (α1^HA^), and β2 γ2S partnering subunits at a 0.5:0.5:1:1 ratio, respectively. Under these conditions, the total α1 expression was significantly reduced compared to WT: 23.6% reduction for I148F (#P < 0.019), 34.4% for R214C (###P < 0.0004), 50.0% for R214H (####P < 0.0001), and 48.5% for A322D (####P < 0.0001; Figure 3A,B). For M263T, T292S, and F325L, total α1 expression was not reduced. In contrast, the WT α1^HA^ allele was expressed at comparable levels in both WT and α1 variant conditions with exception of R214C, which showed 15.9% reduction (###P<.001 vs WT).

**Figure 3.**
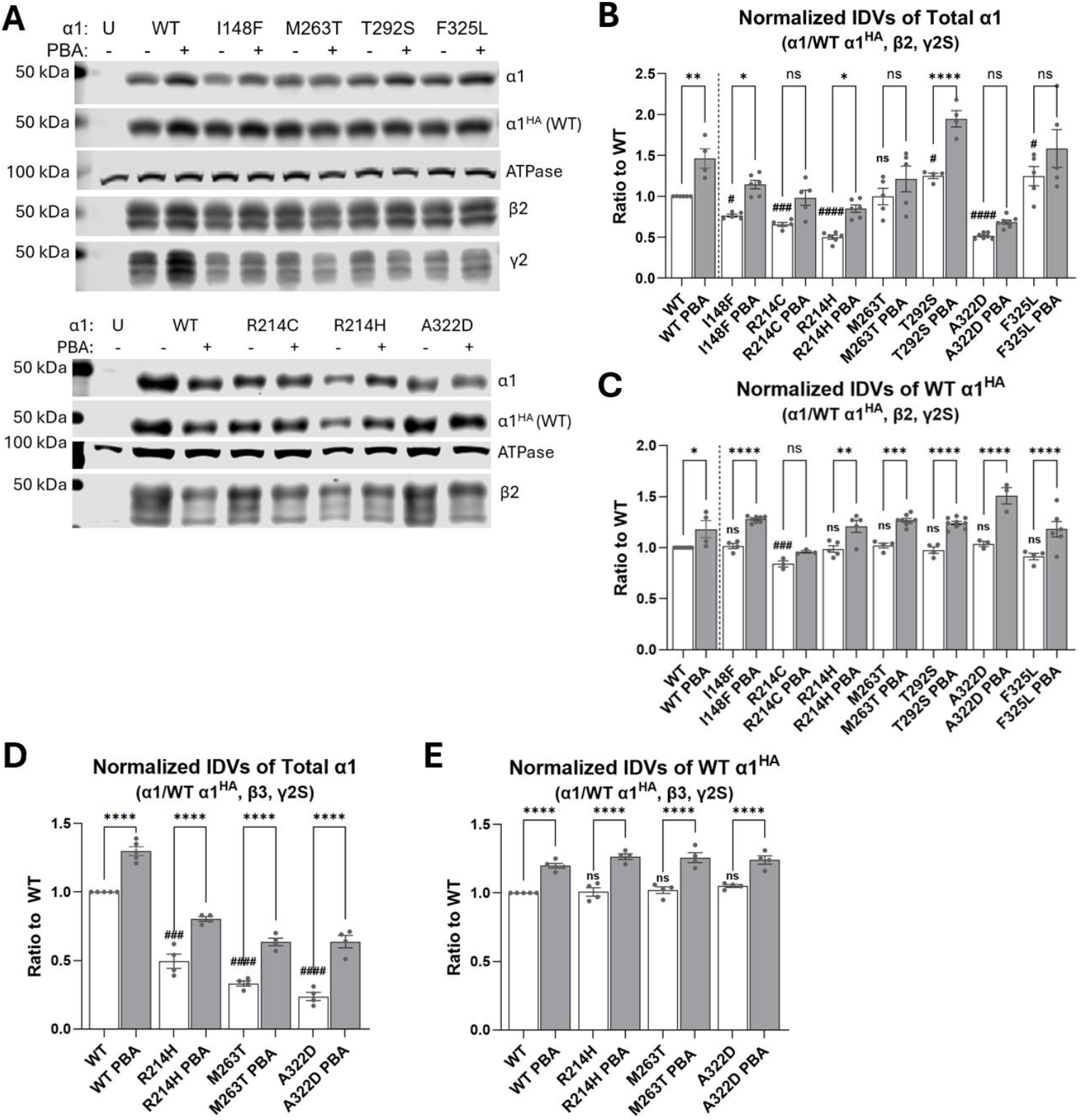
PBA partially restores total and WT α1^HA^ subunit expression in “heterozygous” GABRA1 variant conditions with β2γ2S and β3γ2S receptor assemblies. (**A**) Western blot showing total expression of α1, HA-tagged α1^HA^, β2, and γ2 subunits in HEK293T cells cotransfected with α1, α1^HA^ β2, and γ2S cDNAs (0.5:0.5:1:1 ratio). For each condition, we varied whether the WT or variant α1 was transfected, while α1^HA^ was always kept WT; for PBA treatment, 2mM PBA was added 24 hours prior to harvesting. ATPase was used as a loading control. For a limited number of conditions, we included additional co-expression with the β3 subunit, sample blots for which are shown in Figure S3. (**B**,**C**) Quantification of total α1 (B) and WT α1^HA^ (C) protein levels normalized to untreated WT condition (per blot) in the “heterozygous” α1/α1^HA^β2γ2S conditions. β2 and γ2 quantification is shown in Figure S2. (**D**,**E**) Quantification of total α1 (B) and WT α1^HA^ (C) protein levels normalized to untreated WT condition (per blot) in the subset of variants co-expressed in “heterozygous” α1/α1^HA^β3γ2S conditions. Data are presented as mean ± SEM (n = 4–9; plotted as dot plot on each graph). Statistical analysis was performed using one-way ANOVA followed by Sidak’s multiple comparisons test. Data are presented as mean ± SEM with one-way ANOVA; ####, ###, #P < 0.0001, 0.001, 0.05 untreated variant vs untreated WT by Dunnett’s multiple comparisons test; ****, ***, **, *P < 0.0001, 0.001, 0.01, 0.05 PBA vs corresponding untreated by Sidak’s multiple comparisons test.

### 3.6. PBA treatment increased total and WT α1 expression with “heterozygous” α1α1^HA^β2γ2

Total α1 expression values for the PBA treatment condition were greater than untreated in all cases, but this increase was only statistically significant for WT (46.2% increase; **P < 0.01 vs untreated), I148F (49.7% increase; #P < 0.05 vs untreated), R214H (69.9% increase; ####P < 0.0001 vs untreated), and T292S (55.9% increase; ****P < 0.0001 vs untreated). For I148F and R214H, expression values with treatment surpassed WT baseline expression. Expression values of the WT allele α1^HA^ were also greater than untreated in all cases, and this increase was significant for WT and all variants except R214C: 18.0% for WT (*P < 0.05 vs untreated), 25.8% for I148F (****P < 0.0001 vs untreated), 22.6% for R214H (**P < 0.001 vs untreated), 24.3% for M263T (***P < 0.0001 vs untreated), 27.1% for T292S (****P < 0.0001 vs untreated), and for 29.3% F325L (****P < 0.0001 vs untreated).

### 3.7. Expression of partnering β2 and γ2 subunits with “heterozygous” α1α1^HA^β2γ2

We evaluated γ2 expression for I148F, M263T, T292S, and F325L variants, and observed lower expression compared to WT for all of them: 34.0% decrease for I148F (###P < 0.001 vs untreated WT), 26.5% for M263T (##P < 0.01), 19.5% for T292S (##P < 0.01), and 26.0% for F325L (##P < 0.01). PBA treatment increased γ2 expression in the WT condition by 29.0% (****P < 0.0001 vs untreated); however, we did not observe a similar increase in the variants: I148F showed a moderate 16.7% increase (*P < 0.05); while M263T, T292S, or F325L showed no change nor was there a trend in the values.

Expression of β2 showed no significant changes for the variants, thought the values across the variants were typically slightly lower than WT (Figure S2). Similarly, PBA treatment showed no significant changes in β2 expression across the groups, though values tended to be larger than untreated.

Our sample size and statistical power was lower for γ2 and β2 expression experiments, and it is possible that significant changes in the direction of the indicated trends would be identified with an expanded sample size.

### 3.8. Stronger decrease in total α1 expression for M263T and A322D with β3 partnering subunit

Next, we assessed whether subunit composition influences stability of the variant-containing GABA_A_Rs by performing parallel experiments with the β3 partnering subunit in α1α1^HA^β3γ2S co-expression conditions (Figure 3D–E), which is another highly common synaptic subunit combination in brain. For these experiments, we included only the R214H, M263T, and A322D variants, all of which showed a strong decrease in total α1 expression compared to untreated WT: 50.5% decrease for R214H (####P < 0.0001 vs untreated WT), 66.7% for M263T (####P < 0.0001), and 76.2% for A322D (####P < 0.0001).

While the decrease for R214H was comparable to α1α1^HA^β2γ2S conditions, it was stronger for A322D, and much stronger for M263T, which didn’t show any reduction in α1α1^HA^β2γ2S conditions. Expression of the WT α1^HA^ allele was comparable with WT across these conditions and did not indicate significant dominant negative effects or compensation.

### 3.9 PBA treatment showed stronger increase in total and WT α1 expression with “heterozygous” α1α1^HA^β3γ2

Treatment with PBA strongly increased α1 expression for these variants (Figure 3D): 62.2% for R214H (****P < 0.0001 vs untreated), 91.1% for M263T (****P < 0.0001 vs untreated), and 168.4% for A322D (****P < 0.0001 vs untreated). Again, this was on-par with α1α1^HA^β2γ2S conditions for R214H, but showed a much stronger increase for A322D and M263T. PBA also increased expression of the WT α1^HA^ allele similarly across WT and the three variants (Figure 3E): 25.5% for R214H (****P < 0.0001 vs untreated), 23.3% for M263T (****P < 0.0001 vs untreated), and 18.1% for A322D (***P < 0.001 vs untreated). In summary, the co-expressed subunits seemed to make a palpable difference on stability and expression of variant-containing GABA_A_Rs.

In addition to variants that are the focus of this study, we also evaluated I148P and M263K that have that occur at the same peptide locations as I148F and M263T, respectively (Figure S3). Compared to WT in α1α1^HA^β2γ2S conditions, α1 expression was 63.3% lower for I148P (####P < 0.0001 vs untreated WT) and 77.0% lower for M263K (####P < 0.0001 vs untreated WT); and increased with PBA treatment by 74.4% for I148P (****P < 0.0001 vs untreated) and 165.2% for M263K (****P < 0.0001 vs untreated). Expression of the WT α1^HA^ allele was comparable with WT for I148P and showed a comparable response to PBA treatment (22.1% increase vs untreated; ****P < 0.0001). On the other hand, M263K demonstrated the strongest dominant negative effect we’ve observed among the variants, showing strong decrease in WT α1^HA^ expression (66.2% reduction vs WT; ####P < 0.0001); and PBA treatment was highly effective in treating this phenotype (212.0% increase vs untreated; ****P < 0.0001).

Together, these findings demonstrate that *GABRA1* variants impair total α1 subunit expression under heterozygous conditions, while WT α1 expression is relatively preserved (except M263K). PBA treatment partially restores total α1 levels and increases WT α1 expression across both β2γ2S and β3γ2S receptor assemblies. However, the magnitude of this increase is often smaller than increase seen in total α1 expression, which we also observed with α1_variant_β2γ2S. Therefore, PBA rescue effect appears to involve enhancing the expression of both the WT and variant α1.

### PBA increases α1 expression in a region-specific manner in Gabra1^+/A322D^ mouse brain

Next, we sought to determine whether PBA treatment can improve α1 expression *in vivo*. To do this, we used the *Gabra1*^+/A322D^ mice that have been previously shown to express α1 at about half the WT levels[27]. We followed a 7-day treatment regimen with vehicle or 100 mg/kg/day PBA via intraperitoneal (IP) injection based on our prior studies in *Slc6a1*^+/S295L^ and *Gabrg2*^+/Q390X^ mice[7,9,12]. Then, we collected the mouse brain tissue and quantified α1, β2, and γ2 expression by Western blot on tissue lysates from cortex, cerebellum, hippocampus, and thalamus normalized to ATPase (Figure 4A).

**Figure 4.**
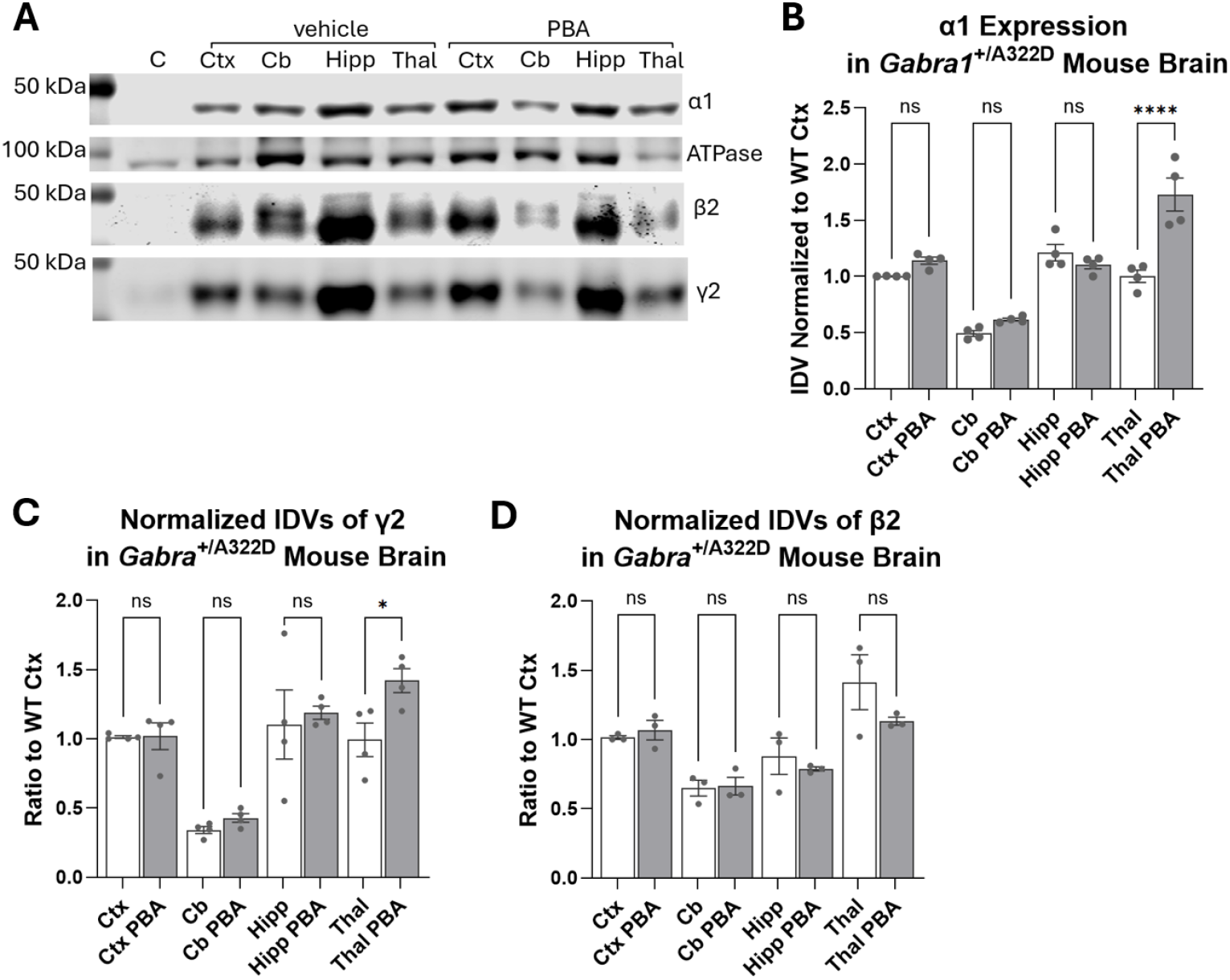
PBA increases α1 subunit expression in a region-specific manner in the brains of *Gabra1*^A322D/−^ mice. (**A**) Western blots showing α1, β2, γ2 subunits expression in tissue lysates from cortex (Ctx), cerebellum (Cb), hippocampus (Hipp), and thalamus (Thal) from *Gabra1*^A322D/−^ mice treated with vehicle or 4-phenylbutyrate (PBA); ATPase was used as a loading control. (**B**) Quantification of α1 protein levels normalized to the vehicle-treated *Gabra1*^A322D/−^ mice. Data are presented as mean ± SEM. n = 4 mice for treated and untreated groups (8 total). ****P < 0.0001 by one-way ANOVA with Sidak’s multiple comparisons test.

We found that PBA treatment significantly increased α1 and γ2 expression in thalamus of *Gabra1*^+/A322D^ mice but did not affect expression in the other regions (Figure 4B,C). In thalamic lysates, α1 expression was 72.8% greater (****P < 0.0001) and γ2 expression was 42.0% greater (*P < 0.05) in PBA group compared to vehicle. On the other hand, we did not detect differences in expression of α1 and γ2 in cortex, cerebellum, and hippocampus; nor in β2 expression across the four regions (Figure 4).

## 4. Discussion

In this study, we evaluated the potential of chemical chaperone and histone deacetylate inhibitor PBA to treat a cellular pathology in GABA_A_R expression for seven *GABRA1* variants I148F, R214C, R214H, M253T, T292S, A322D, and F325L associated with neurologic disease in human patients. Using *in silico* modeling, we identified that the variants occur in highly conserved residues and act to destabilize the α1 subunit to varying degrees. The variants demonstrated reduced total (except F325L) and surface expression in HEK293T cells. For half of the variants, the reduction in total α1 expression persisted in “heterozygous” co-expression with WT α1 α1^HA^, though dominant negative effects on the WT subunit were not observed (except R214C). PBA treatment significantly improved both total (except R214C) and surface expression of the variants; and WT α1 subunit in the control condition and with “heterozygous” co-expression with variants. These findings suggest a shared folding and trafficking pathology of varying severity for these variants; and showcase that PBA is an effective treatment that improves expression of both mutant and wildtype α1 alleles.

At the structural level, computational modeling predicted that all variants exhibit destabilizing effects on the α1 subunit, consistent with disruption of intermolecular interactions within the N-terminal domain and transmembrane helices. Such perturbations are expected to impair subunit folding and assembly, thereby compromising receptor maturation. This is consistent with the negative ΔΔG predicted change in stability values across multiple computational platforms. The ΔΔG values also matched our predictions based on structural visualization of the mutations and the highly conserved status of the residues. However, ΔΔG magnitude was a weak predictor for magnitude of decrease in subunit expression in our experiments. For example, the F325L variant showed the least change in α1 expression among the variants, which matched the smaller ΔΔG magnitude relative to other variants. Contrary to this trend, however, the A322D variant exhibited the strongest decrease in α1 expression but displayed the smallest predicted ΔΔG magnitude.

We found that most variants showed a similarly strong decrease in α1 expression when expressed as the only α subunit; but showed differences when co-expressed with WT α1^HA^. In α1^variant^β2γ2 conditions, both total and surface α1 expression decreased in the 61-85% range compared to WT control with M263T the least and A322D the most affected; which corresponded with a decrease in current amplitude of GABA-evoked currents. F325L was the only exception and only showed a decrease in surface expression. However, in “heterozygous” α1^variant^α1^HA^β2γ2 conditions, only the variants I148F, R214C, R214H, and A322D showed decreased α1 expression, while the M263T and T292S variants did not. This was unlikely to happen from compensatory WT α1 expression alone because WT α1^HA^ expression levels were comparable between WT and most variants. One possible explanation is that the WT α1^HA^ subunit stabilized formation of GABA_A_Rs containing a single WT and a single variant α1 subunit, which has previously been reported for specific configurations with the A322D variant[15]. This stabilizing effect would differ by variant, and is particularly plausible in case of M263T based on the milder expression phenotype under α1^variant^β2γ2 conditions. This could also explain why M263T variant showed a significant drop in α1 expression in the α1^variant^α1^HA^β3γ2 condition – substitution of β3 for β2 subunit appeared to form a less stable configuration with the variant.

We did not observe significant dominant negative effects on the WT α1^HA^ subunit with exception of the R214C variant. However, expression of the partnering subunits β2 and γ2 was decreased when co-expressed with the mutant α1 variants; this was particularly consistent for γ2 including both α1^variant^β2γ2 and α1^variant^α1^HA^β2γ2 conditions. In cases when no statistically significant decrease was observed, we often still observed lower expression values than WT. Whether this constitutes a dominant negative effect is debatable, however, since in these conditions presence of a functional α1 subunit is necessary for receptor formation. Therefore, it is also likely that this decrease was associated with reduced availability of partnering α1.

Compared to existing expression studies, our findings match some of the reports but also show some differences. Our findings match the extensive prior A322D characterization including reduced total and surface α1 expression, a mild dominant negative effect, and a strong reduction in GABA_A_R current magnitude[14-16]; and the initial characterization of R214C showing reduced receptor current with decreased α1 membrane and total expression[20,21]. Our findings also match reduced GABA_A_R current magnitude reported by Hernandez et al for R214C and R214H; unlike the authors, we observed reduced α1 expression, while the authors argued for primarily a gating defect[19]. Our findings for T292S differed from a report by Chen et al of gain in GABA_A_R function associated with increased ligand sensitivity but normal total and surface expression [24]. While we did not find evidence contrary to changes in GABA_A_R sensitivity, T292S showed a significant decrease in both total and surface α1 expression in our study, and only showed normal levels when co-expressed with WT α1^HA^. We are unsure as to the source of discrepancy in our expression results, as the aforementioned studies[19,24] use similar methods including the HEK293T cell system and the β2 and γ2 partnering subunits, and do not use tethered constructs that could have an artificial stabilizing effect. For the I148F, F325L, and M253T variants, our study is the first functional characterization.

Treatment with PBA caused a significant increase in expression of α1 protein including WT and mutant variants. For WT α1, we observed higher total and surface α1 expression in WT control α1β2γ2 conditions and α1α1^HA^β2γ2 conditions. Additionally, WT α1^HA^ was expressed at higher levels in “heterozygous” α1α1^HA^β2γ2 co-expression with most α1 variants, and the magnitude of this increase was generally comparable with WT α1 control. Therefore, it seems that one effect of PBA was to increase α1 expression regardless of presence of the mutant protein, which is beneficial in case of a haploinsufficiency disorder. On the other hand, PBA also increased α1 expression in cases when only the variant α1 was transfected, indicating that the variant α1 proteins were also stabilized and increased in expression. Therefore, increased total α1 expression in co-expression conditions likely reflects improvements in both the WT and mutant α1 protein expression as well as general stabilization of the GABA_A_R pentamer.

The R214C variant was an exception and showed the least improvement from PBA treatment. PBA treatment increased surface α1 expression for R214C in α1β2γ2 conditions on par with other variants; but total α1 expression was unchanged. Additionally, R214C appeared to have a dominant negative effect on WT α1-HA in “heterozygous” α1α1^HA^β2γ2 conditions, showing reduced α1^HA^ expression compared to WT and not showing an α1^HA^ increase with PBA treatment. For a future study, it would be helpful to evaluate how PBA treatment affects the surface expression of α1-HA when co-expressed with R214C variant.

We showed that PBA treatment also improved α1 expression *in vivo* in *Gabra1*^+/A322D^ mice. *Gabra1*^+/A322D^ mice express roughly half the WT levels of α1[27], and we observed a 72.8% increase in thalamic α1 expression with PBA treatment. However, we did not observe an increase in the other three regions, though α1 is expressed widely throughout the brain with high expression in all of the studied regions[33]. This indicates a regional effect and reflects increased *in vivo* complexity compared to *in vitro* reductionism, including presence of many other partnering subunits. A rescue in these regions may be better evaluated by surface expression and synaptic function. Nonetheless, we found these results promising, and believe there is high potential in evaluating whether PBA treatment can reduce the seizure burden in *Gabra1*^+/A322D^ mice[27] as it has for other models[7,9].

Together, our findings support a model of a common proteostasis pathology for *GABRA1* variants that is highly responsive to PBA therapy (Figure 5). In our prior work, we demonstrated this for neurologic disorders associated with *SLC6A1* and *GABRG2* variants. Mutations destabilize the variant proteins, causing defective folding, inefficient assembly, and reduced trafficking. Misfolded or incompletely assembled subunits are likely to be retained within the endoplasmic reticulum and subjected to degradation pathways, ultimately reducing receptor surface availability (Fu et al., 2022; Williams et al., 2025). The significant and consistent reduction in α1 expression and restorative benefits of PBA therapy that we observed for most *GABRA1* variants are evidence for this model, and strengthen the rationale of clinical trial development for *GABRA1* disorders. Moreover, more targeted future evaluations would be beneficial in further testing this model, including a direct evaluation of variant α1 subunit trafficking in endoplasmic reticulum; interactions between variant α1 subunits and endoplasmic reticulum proteostasis components including chaperones BiP and calnexin; thorough electrophysiologic evaluation of the variants and PBA therapy effects; and effects of PBA on seizure burden in mouse models including *Gabra1*^+/A322D^ mice.

**Figure 5.**
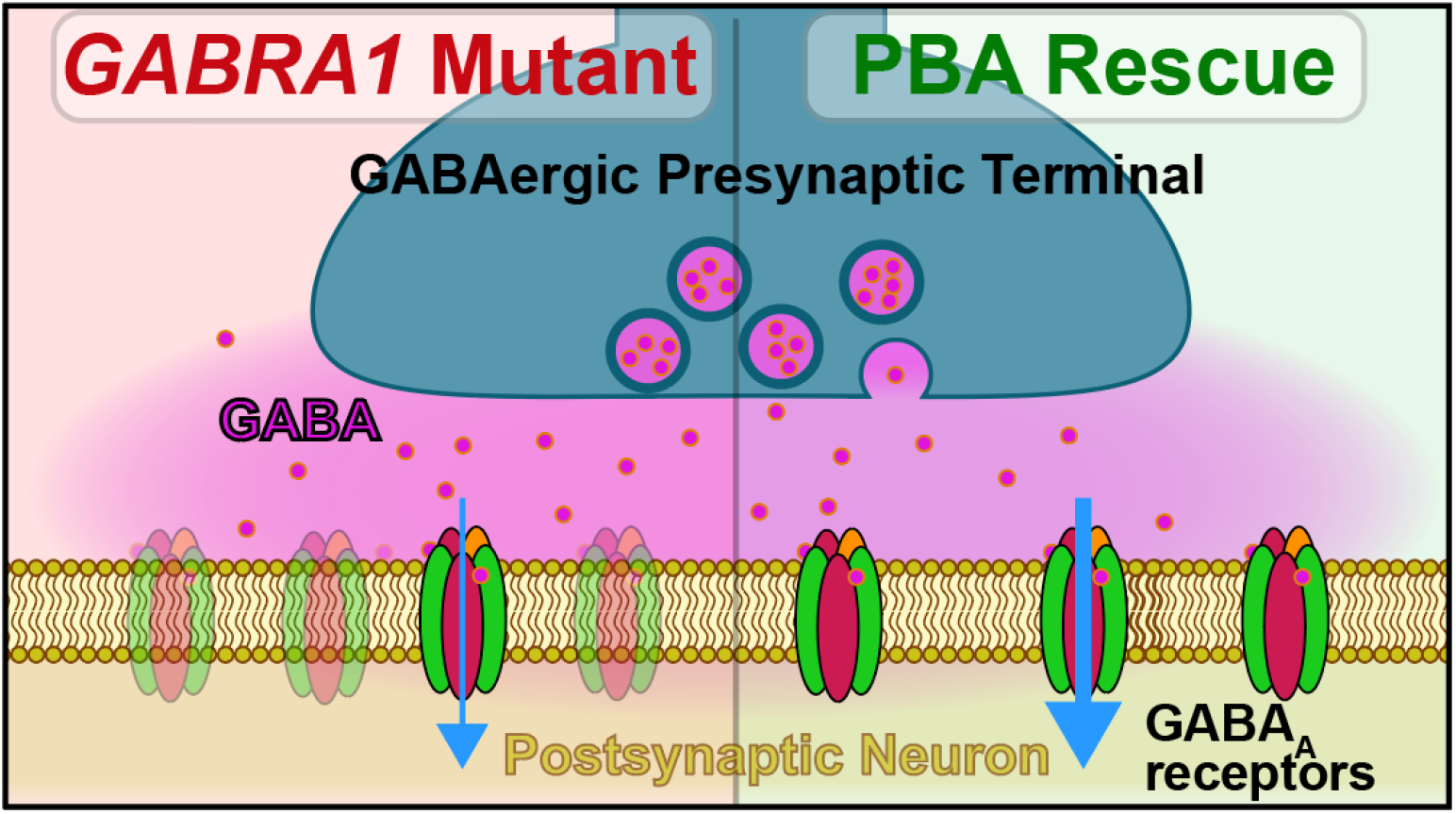
Our proposed model of a shared patho-mechanism for many *GABRA1* disorders, based on this study and previous work in similar disorders. *GABRA1* variants affect GABA_A_R proteostasis, reducing the receptor number at the synapse and weaking synaptic responses. PBA rescue stabilized proteostasis and increases GABA_A_R subunit expression, improving receptor number at the synapse and restoring synaptic responses.

## Supplementary Materials

The following supporting information can be downloaded: Supplemental Figures.docx including Figures S1-S3.

## Author Contributions

Conceptualization, JQK; methodology, JKQ; investigation (electrophysiology) LM; investigation (biochemistry) ZDS, WS, MBD, and GXH, RSE; investigation (flow cytometry) ZDS; investigation (alignment), KZ; investigation (modeling), ZDS; formal analysis, ZDS and KZ; resources (patient), SP and RC; writing—original draft preparation, ZDS and KZ; writing— review and editing, KZ and ZDS; visualization, ZDS and KZ; supervision, JQK; project administration, JQK; funding acquisition, JQK. All authors have read and agreed to the published version of the manuscript.

## Institutional Review Board Statement

All animal procedures were performed in accordance with protocols and were approved by the Vanderbilt University Institutional Animal Care and Use Committee (IACUC).

## Data Availability Statement

Data is available upon request from the corresponding author.

## Acknowledgments

We would like to acknowledge Monica Joanna Elnekaveh and Agustina Fernández Giambruno from CURE GABA-A Variants for providing the GABRA1 variants included in this study. We would like to acknowledge Sarah Poliquin for their contribution in providing patient resources. We would also like to acknowledge Dr. Martin Gallagher for providing the *Gabra*^+/A322D^ mice. We would also like to acknowledge Dr. Juexin Wang for guidance in AI structural Modeling.

## References

1. Feng, Y.;Wei, Z.H.;Liu, C.;Li, G.Y.;Qiao, X.Z.;Gan, Y.J.;Zhang, C.C.;Deng, Y.C. Genetic variations in GABA metabolism and epilepsy. Seizure 2022, 101, 22–29, doi:10.1016/j.seizure.2022.07.007.

2. Kang, J.Q. Defects at the crossroads of GABAergic signaling in generalized genetic epilepsies. Epilepsy Res 2017, 137, 9–18, doi:10.1016/j.eplepsyres.2017.08.013.

3. Fu, X.;Wang, Y.J.;Kang, J.Q.;Mu, T.W. GABA(A) Receptor Variants in Epilepsy. In Epilepsy, Czuczwar, S.J., Ed.; Brisbane (AU), 2022.

4. Stone, A.;Burre, J.;Wayland, N.;Grinspan, Z.M. Phenylbutyrate for monogenetic epilepsy: Literature review. Epilepsy Res 2025, 217, 107621, doi:10.1016/j.eplepsyres.2025.107621.

5. Sugiyama, T.;Nishitoh, H. Neurodegenerative diseases associated with the disruption of proteostasis and their therapeutic strategies using chemical chaperones. J Biochem 2024, 176, 179–186, doi:10.1093/jb/mvae048.

6. Kang, J.Q.;Shen, W.;Zhou, C.;Xu, D.;Macdonald, R.L. The human epilepsy mutation GABRG2(Q390X) causes chronic subunit accumulation and neurodegeneration. Nat Neurosci 2015, 18, 988–996, doi:10.1038/nn.4024.

7. Shen, W.;Flamm, C.;Delahanty, A.J.;Casteel, E.;Biven, M.;DeLeeuw, M.B.;Poliquin, S.;Nwosu, G.;Randhave, K.;Kang, J.Q. 4-Phenylbutyrate promoted wild-type gamma-aminobutyric acid type A receptor trafficking, reduced endoplasmic reticulum stress, and mitigated seizures in Gabrg2(+/Q390X) mice associated with Dravet syndrome. Epilepsia 2023, doi:10.1111/epi.17779.

8. Mermer, F.;Poliquin, S.;Rigsby, K.;Rastogi, A.;Shen, W.;Romero-Morales, A.;Nwosu, G.;McGrath, P.;Demerast, S.;Aoto, J.;et al. Common molecular mechanisms of SLC6A1 variant-mediated neurodevelopmental disorders in astrocytes and neurons. Brain 2021, 144, 2499–2512, doi:10.1093/brain/awab207.

9. Nwosu, G.;Mermer, F.;Flamm, C.;Poliquin, S.;Shen, W.;Rigsby, K.;Kang, J.Q. 4-Phenylbutyrate restored gammaaminobutyric acid uptake and reduced seizures in SLC6A1 patient variant-bearing cell and mouse models. Brain Commun 2022, 4, fcac144. doi:10.1093/braincomms/fcac144.

10. Shen, W.;Nwosu, G.;Honer, M.;Clasadonte, J.;Schmalzbauer, S.;Biven, M.;Langer, K.;Flamm, C.;Poliquin, S.;Mermer, F.;et al. gamma-Aminobutyric acid transporter and GABA(A) receptor mechanisms in Slc6a1(+/A288V) and Slc6a1(+/S295L) mice associated with developmental and epileptic encephalopathies. Brain Commun 2024, 6, fcae110. doi:10.1093/braincomms/fcae110.

11. DeLeeuw, M.B.;Shen, W.;Tian, X.;Ding, C.;Randhave, K.;Kang, J.Q. 4-Phenylbutyrate restored GABA uptake, mitigated seizures in SLC6A1 and SLC6A11 microdeletions/3p-syndrome: From cellular models to human patients. Epilepsy Res 2025, 210, 107514, doi:10.1016/j.eplepsyres.2025.107514.

12. Randhave, K.;Zavalin, K.;Kang, J.Q. 4-Phenylbutyrate rescues neurobehavioral phenotypes in SLC6A1-related encephalopathy. Epilepsy Res 2025, 220, 107708, doi:10.1016/j.eplepsyres.2025.107708.

13. Grinspan, Z.M.;Burré, J.;Cross, J.;Ross, M.E.;Stone, A.;Basma, N.;Gao, K.;Kang, J.-Q.;Lim, J.;Dubow, E.;et al. 4-Phenylbutyrate for STXBP1 and SLC6A1. Safety, tolerability, seizure, and EEG outcomes. A case series at 2 centers. medRxiv 2024, doi:10.1101/2024.11.06.24316676.

14. Cossette, P.;Liu, L.;Brisebois, K.;Dong, H.;Lortie, A.;Vanasse, M.;Saint-Hilaire, J.M.;Carmant, L.;Verner, A.;Lu, W.Y.;et al. Mutation of GABRA1 in an autosomal dominant form of juvenile myoclonic epilepsy. Nat Genet 2002, 31, 184–189, doi:10.1038/ng885.

15. Gallagher, M.J.;Song, L.;Arain, F.;Macdonald, R.L. The juvenile myoclonic epilepsy GABA(A) receptor alpha1 subunit mutation A322D produces asymmetrical, subunit position-dependent reduction of heterozygous receptor currents and alpha1 subunit protein expression. J Neurosci 2004, 24, 5570–5578, doi:10.1523/JNEUROSCI.1301-04.2004.

16. Krampfl, K.;Maljevic, S.;Cossette, P.;Ziegler, E.;Rouleau, G.A.;Lerche, H.;Bufler, J. Molecular analysis of the A322D mutation in the GABA receptor alpha-subunit causing juvenile myoclonic epilepsy. Eur J Neurosci 2005, 22, 10–20, doi:10.1111/j.1460-9568.2005.04168.x.

17. Ding, L.;Feng, H.J.;Macdonald, R.L.;Botzolakis, E.J.;Hu, N.;Gallagher, M.J. GABA(A) receptor alpha1 subunit mutation A322D associated with autosomal dominant juvenile myoclonic epilepsy reduces the expression and alters the composition of wild type GABA(A) receptors. J Biol Chem 2010, 285, 26390–26405, doi:10.1074/jbc.M110.142299.

18. Gallagher, M.J.;Ding, L.;Maheshwari, A.;Macdonald, R.L. The GABAA receptor alpha1 subunit epilepsy mutation A322D inhibits transmembrane helix formation and causes proteasomal degradation. Proc Natl Acad Sci U S A 2007, 104, 12999–13004, doi:10.1073/pnas.0700163104.

19. Hernandez, C.C.;Tian, X.;Hu, N.;Shen, W.;Catron, M.A.;Yang, Y.;Chen, J.;Jiang, Y.;Zhang, Y.;Macdonald, R.L. Dravet syndrome-associated mutations in GABRA1, GABRB2 and GABRG2 define the genetic landscape of defects of GABA(A) receptors. Brain Commun 2021, 3, fcab033. doi:10.1093/braincomms/fcab033.

20. Bai, Y.F.;Chiu, M.;Chan, E.S.;Axerio-Cilies, P.;Lu, J.;Huh, L.;Connolly, M.B.;Guella, I.;Farrer, M.J.;Xu, Z.D.;et al. Pathophysiology of and therapeutic options for a GABRA1 variant linked to epileptic encephalopathy. Mol Brain 2019, 12, 92, doi:10.1186/s13041-019-0513-9.

21. Bai, Y.F.;Chiu, M.;Chan, E.S.;Axerio-Cilies, P.;Lu, J.;Huh, L.;Connolly, M.B.;Guella, I.;Farrer, M.J.;Xu, Z.D.;et al. Correction to: Pathophysiology of and therapeutic options for a GABRA1 variant linked to epileptic encephalopathy. Mol Brain 2020, 13, 50, doi:10.1186/s13041-020-00593-6.

22. Johannesen, K.;Marini, C.;Pfeffer, S.;Moller, R.S.;Dorn, T.;Niturad, C.E.;Gardella, E.;Weber, Y.;Sondergard, M.;Hjalgrim, H.;et al. Phenotypic spectrum of GABRA1: From generalized epilepsies to severe epileptic encephalopathies. Neurology 2016, 87, 1140–1151, doi:10.1212/WNL.0000000000003087.

23. Zhang, L.;Liu, X. Clinical phenotype and genotype of children with GABA(A) receptor alpha1 subunit gene-related epilepsy. Front Neurol 2022, 13, 941054, doi:10.3389/fneur.2022.941054.

24. Chen, W.;Ge, Y.;Lu, J.;Melo, J.;So, Y.W.;Juneja, R.;Liu, L.;Wang, Y.T. Distinct Functional Alterations and Therapeutic Options of Two Pathological De Novo Variants of the T292 Residue of GABRA1 Identified in Children with Epileptic Encephalopathy and Neurodevelopmental Disorders. Int J Mol Sci 2022, 23, doi:10.3390/ijms23052723.

25. Demos, M.;Guella, I.;DeGuzman, C.;McKenzie, M.B.;Buerki, S.E.;Evans, D.M.;Toyota, E.B.;Boelman, C.;Huh, L.L.;Datta, A.;et al. Diagnostic Yield and Treatment Impact of Targeted Exome Sequencing in Early-Onset Epilepsy. Front Neurol 2019, 10, 434, doi:10.3389/fneur.2019.00434.

26. Kodera, H.;Ohba, C.;Kato, M.;Maeda, T.;Araki, K.;Tajima, D.;Matsuo, M.;Hino-Fukuyo, N.;Kohashi, K.;Ishiyama, A.;et al. De novo GABRA1 mutations in Ohtahara and West syndromes. Epilepsia 2016, 57, 566–573, doi:10.1111/epi.13344.

27. Arain, F.;Zhou, C.;Ding, L.;Zaidi, S.;Gallagher, M.J. The developmental evolution of the seizure phenotype and cortical inhibition in mouse models of juvenile myoclonic epilepsy. Neurobiol Dis 2015, 82, 164–175, doi:10.1016/j.nbd.2015.05.016.

28. Wang, J.;Shen, D.;Xia, G.;Shen, W.;Macdonald, R.L.;Xu, D.;Kang, J.Q. Differential protein structural disturbances and suppression of assembly partners produced by nonsense GABRG2 epilepsy mutations: implications for disease phenotypic heterogeneity. Sci Rep 2016, 6, 35294, doi:10.1038/srep35294.

29. Kang, J.Q.;Shen, W.;Macdonald, R.L. Two molecular pathways (NMD and ERAD) contribute to a genetic epilepsy associated with the GABA(A) receptor GABRA1 PTC mutation, 975delC, S326fs328X. J Neurosci 2009, 29, 2833–2844, doi:10.1523/JNEUROSCI.4512-08.2009.

30. Poliquin, S.;Nwosu, G.;Randhave, K.;Shen, W.;Flamm, C.;Kang, J.Q. Modulating Endoplasmic Reticulum Chaperones and Mutant Protein Degradation in GABRG2(Q390X) Associated with Genetic Epilepsy with Febrile Seizures Plus and Dravet Syndrome. Int J Mol Sci 2024, 25, doi:10.3390/ijms25094601.

31. Nwosu, G.I.;Shen, W.;Zavalin, K.;Poliquin, S.;Randhave, K.;Flamm, C.;Biven, M.;Langer, K.;Kang, J.Q. GABA(A) Receptor beta3 Subunit Mutation N328D Heterozygous Knock-in Mice Have Lennox-Gastaut Syndrome. Int J Mol Sci 2023, 24, doi:10.3390/ijms24098458.

32. Cheng, W.W.L.;Arcario, M.J. Structure-Function Studies of Pentameric Ligand-Gated Ion Channels: Combining Experimental and Computational Approaches. Adv Exp Med Biol 2026, 1497, 287–306, doi:10.1007/978-3-032-07523-9_11.

33. Hortnagl, H.;Tasan, R.O.;Wieselthaler, A.;Kirchmair, E.;Sieghart, W.;Sperk, G. Patterns of mRNA and protein expression for 12 GABAA receptor subunits in the mouse brain. Neuroscience 2013, 236, 345–372, doi:10.1016/j.neuroscience.2013.01.008.

