## Supplemental Figures for "4-Phenylbutyrate Rescue in *GABRA1* Variants Associated with Developmental Epileptic Encephalopathies: from Cell and Mouse Models to Human"

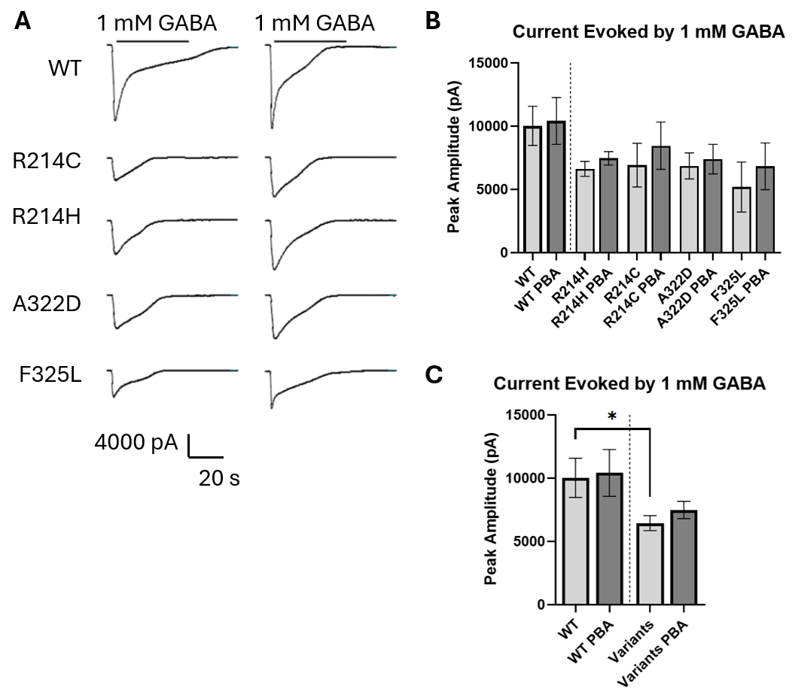
 Figure S1. Lower peak GABA current in HEK293T cells expressing α1-variant GABA_A_Rs. Current was elicited by 1 mM GABA application in HEK293T cells expressing α1β2γ2 at 1:1:0.5 ratio with and without PBA treatment; either WT or variant α1 subunits were expressed. (A) Sample traces. (B) Quantification of peak current amplitude for all variants. (C) Quantification of peak current amplitude with variants pooled for higher statistical power. n = 10 cells for WT, 8 for WT + PBA, 19 for Variants, 19 for Variants + PBA, 6 for R214H, 4 for R214H + PBA, 4 for R214C, 4 for R214C + PBA, 5 for A322D, 6 for A322D + PBA, 4 for F325L, and 5 for F325L + PBA. Data shown as mean ± SEM with one-way ANOVA; *P < 0.05 by Sidak’s multiple comparisons.

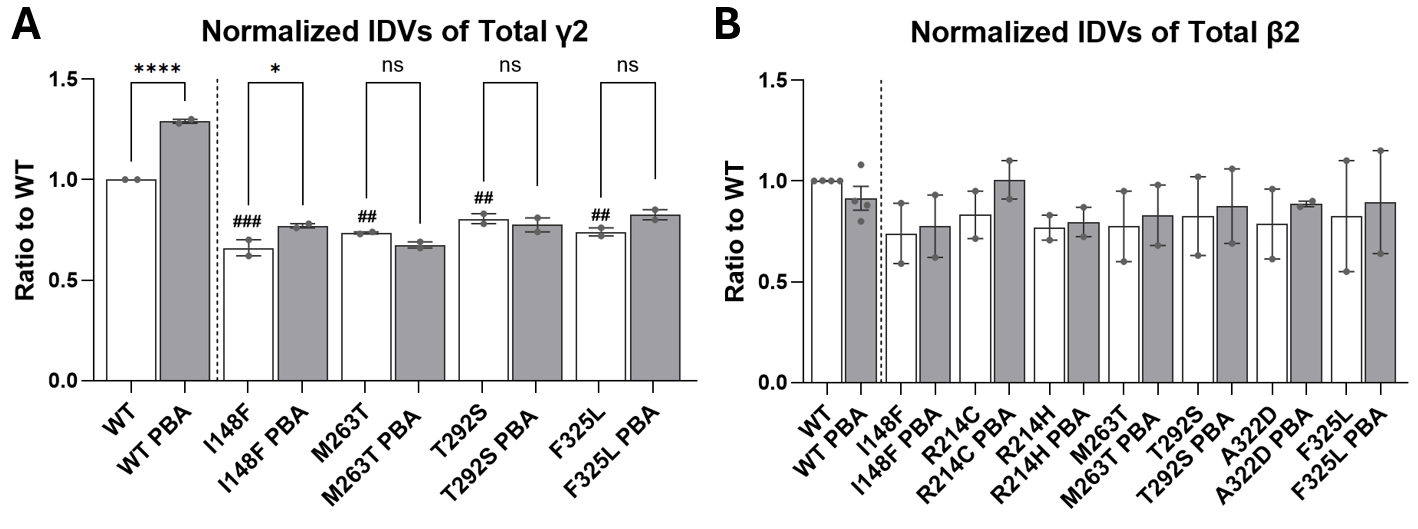
 Figure S2. Expression of β2 and γ2 partnering subunits in “heterozygous” co-expression conditions with and without PBA treatment. Graphs display (A) γ2 and (B) β2 expression levels measured by Western blot in HEK293T cells co-expressing WT or variant α1, WT α1^HA^, β2, and γ2S partnering subunits at a 0.5:0.5:1:1 ratio. These data match Western blot images and α1 quantification shown in Figure 3A-C. Graphs show mean ± SEM with one-way ANOVA with Sidak multiple comparisons; ###, ##P < 0.001, 0.01 vs untreated WT; ****,*P < 0.0001, 0.05 vs respective untreated condition as indicated by brackets.

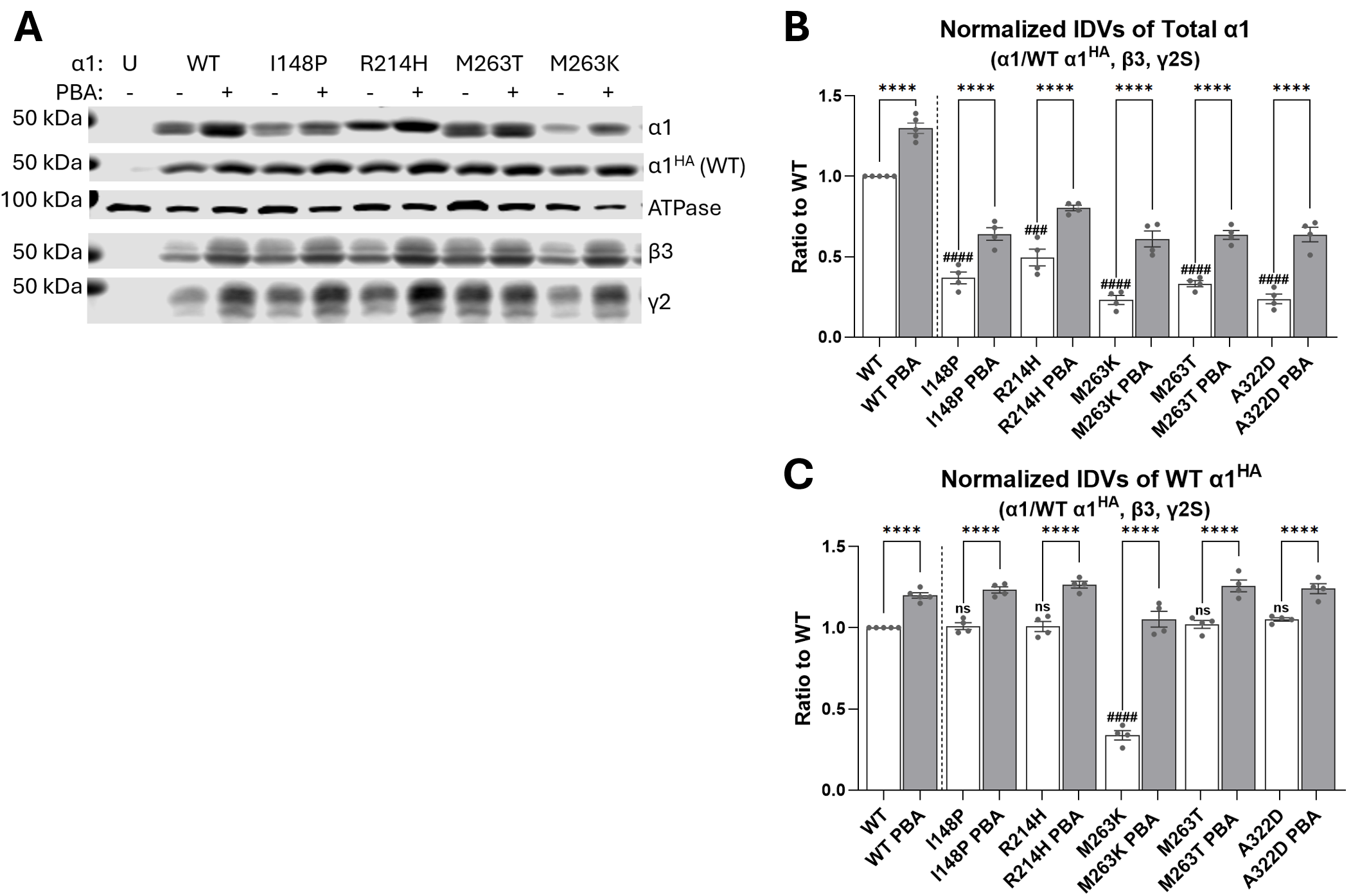
 Figure S3. Total α1 and WT α1^HA^ expression with β3 and γ2 partnering subunits in “heterozygous” co-expression conditions with and without PBA treatment (expanded from Figure 3D-E). (A) Sample Western blot showing the expressed α1, α1^HA^, β3, and γ2S subunits for some of the subunit conditions. (B) Quantification of total α1 expression. (C) Quantification of total WT α1^HA^ expression. Note that only the R214H, M263T, and A322D variants out of the main dataset in this study were tested. Additional variants I148P and M263K that share the residue location with I148F and M263T variants investigated in this study were included. Graphs show mean ± SEM with one-way ANOVA; ####, ###P < 0.0001, 0.001 vs untreated WT by Dunnett’s multiple comparisons; ****P < 0.0001 vs respective untreated condition as indicated by brackets by Sidak’s multiple comparisons.
